## Supplementary material revised for "H pilin cyclisation and pilus biogenesis are promiscuous but electrostatic perturbations impair conjugation efficiency"

**A**

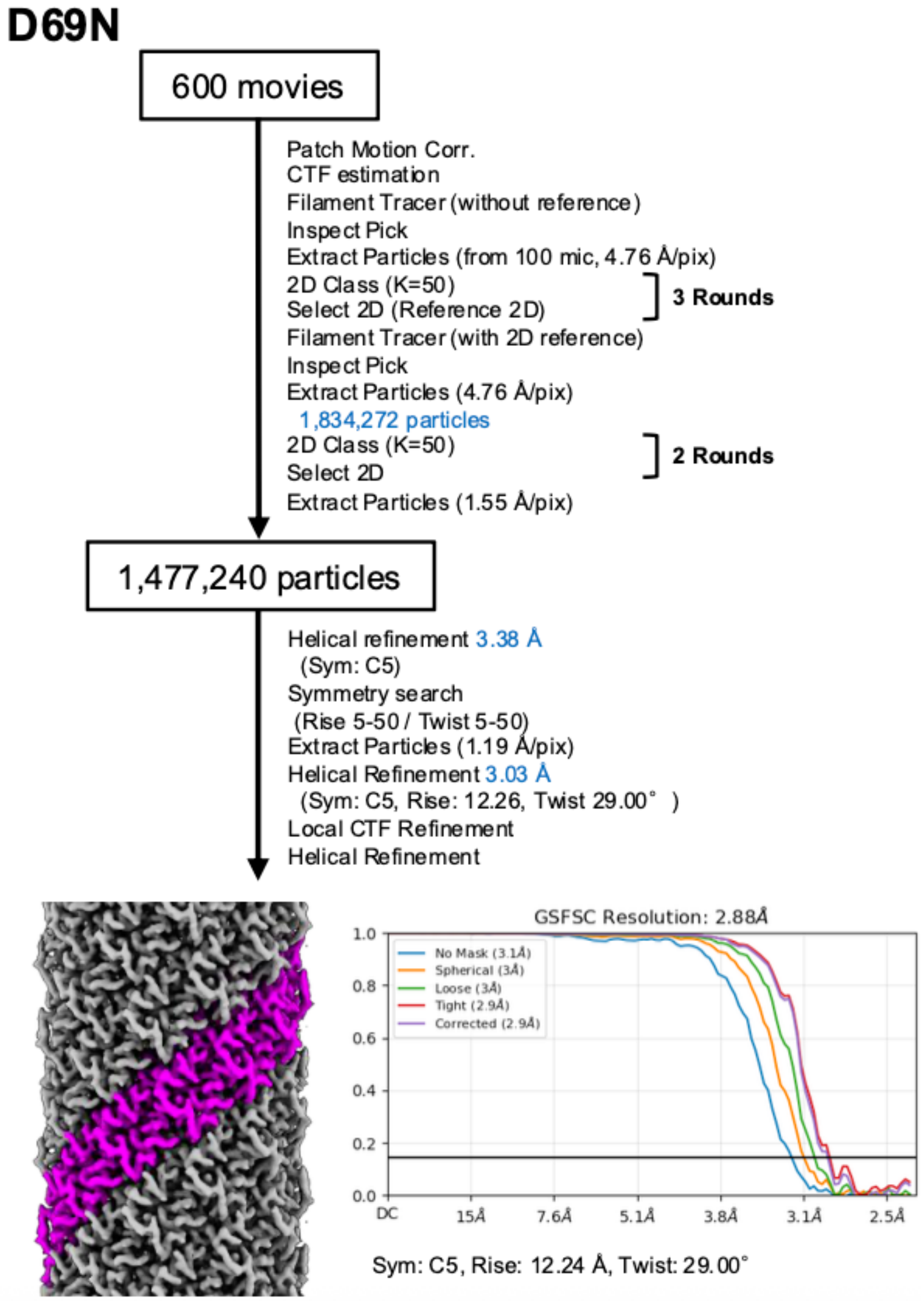
D69N
600 movies
Patch Motion Corr.
CTF estimation
Filament Tracer (without reference)
Inspect Pick
Extract Particles (from 100 mic, 4.76 Å/pix)
2D Class (K=50)
Select 2D (Reference 2D)
3 Rounds
Filament Tracer (with 2D reference)
Inspect Pick
Extract Particles (4.76 Å/pix)
1,834,272 particles
2D Class (K=50)
Select 2D
2 Rounds
Extract Particles (1.55 Å/pix)
1,477,240 particles
Helical refinement 3.38 Å
(Sym: C5)
Symmetry search
(Rise 5-50 / Twist 5-50)
Extract Particles (1.19 Å/pix)
Helical Refinement 3.03 Å
(Sym: C5, Rise: 12.26, Twist 29.00° )
Local CTF Refinement
Helical Refinement
GSFSC Resolution: 2.88Å
No Mask (3.1Å)
Spherical (3Å)
Loose (3Å)
Tight (2.9Å)
Corrected (2.9Å)
1.0
0.8
0.6
0.4
0.2
0.0
DC
15Å
7.6Å
5.1Å
3.8Å
3.1Å
2.5Å
Sym: C5, Rise: 12.24 Å, Twist: 29.00°

**B**

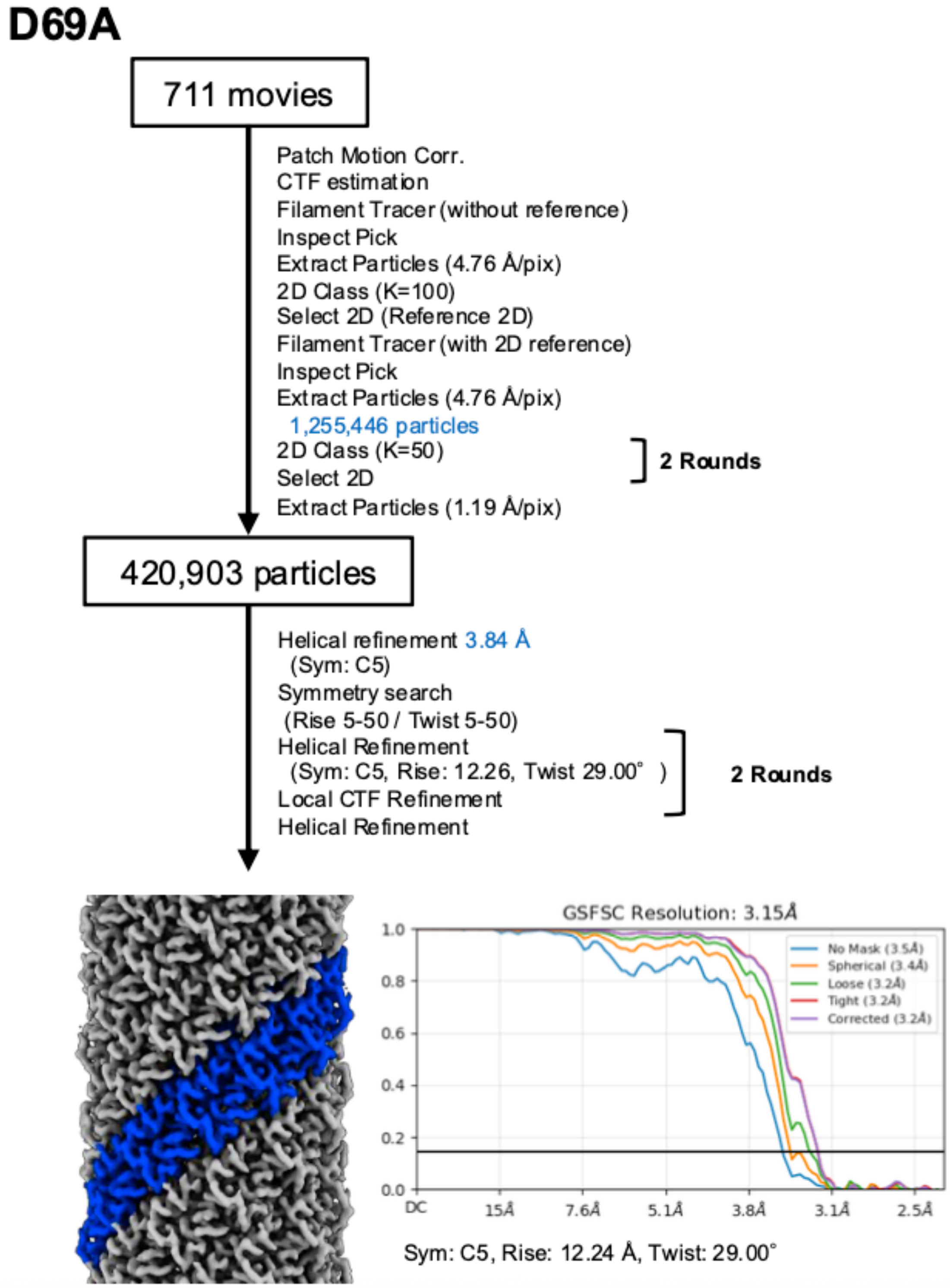
D69A
711 movies
Patch Motion Corr.
CTF estimation
Filament Tracer (without reference)
Inspect Pick
Extract Particles (4.76 Å/pix)
2D Class (K=100)
Select 2D (Reference 2D)
Filament Tracer (with 2D reference)
Inspect Pick
Extract Particles (4.76 Å/pix)
1,255,446 particles
2D Class (K=50)
Select 2D
2 Rounds
Extract Particles (1.19 Å/pix)
420,903 particles
Helical refinement 3.84 Å
(Sym: C5)
Symmetry search
(Rise 5-50 / Twist 5-50)
Helical Refinement
(Sym: C5, Rise: 12.26, Twist 29.00° )
Local CTF Refinement
2 Rounds
Helical Refinement
GSFSC Resolution: 3.15Å
No Mask (3.5Å)
Spherical (3.4Å)
Loose (3.2Å)
Tight (3.2Å)
Corrected (3.2Å)
1.0
0.8
0.6
0.4
0.2
0.0
DC
15Å
7.6Å
5.1Å
3.8Å
3.1Å
2.5Å
Sym: C5, Rise: 12.24 Å, Twist: 29.00°

**C**

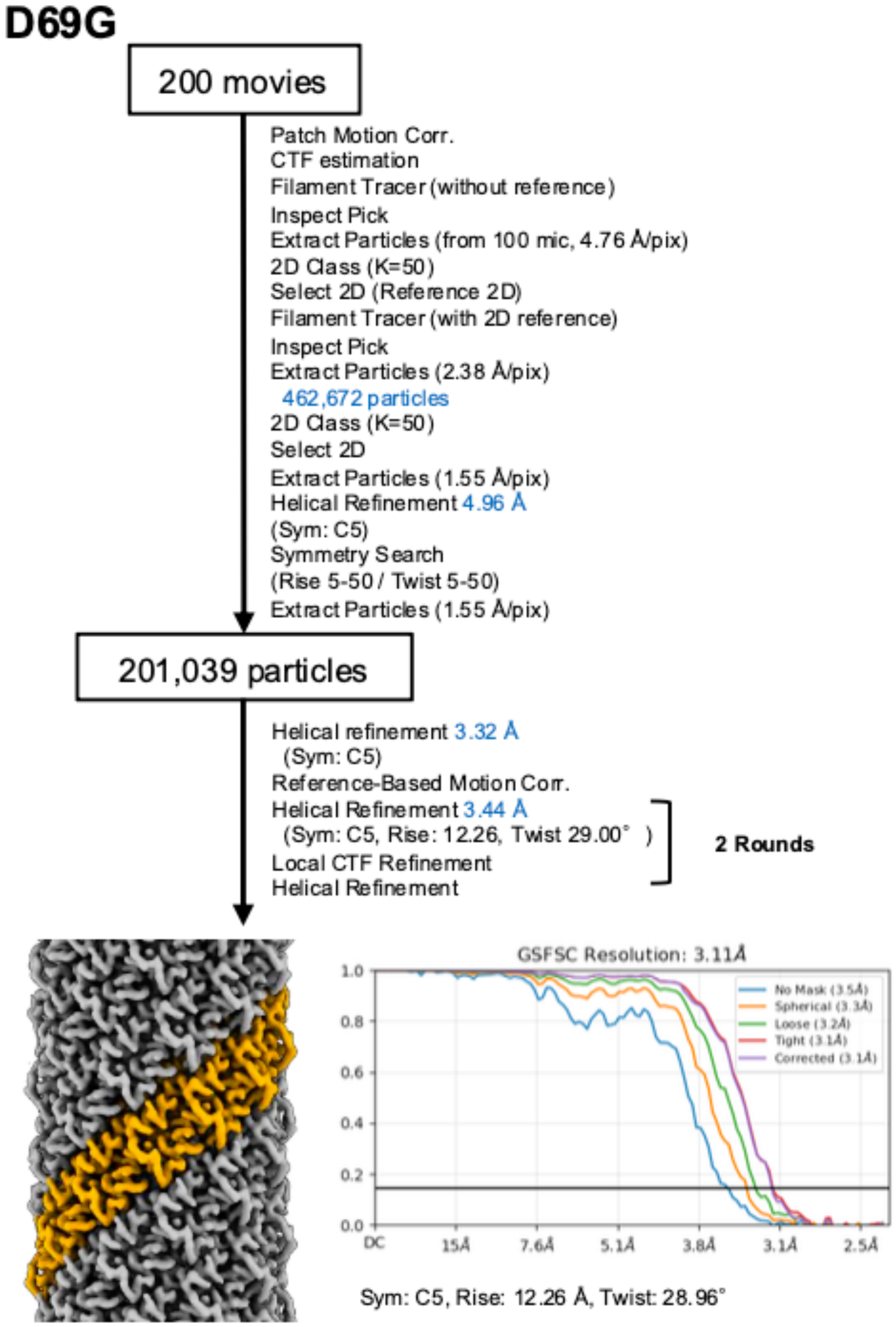
D69G
200 movies
Patch Motion Corr.
CTF estimation
Filament Tracer (without reference)
Inspect Pick
Extract Particles (from 100 mic, 4.76 Å/pix)
2D Class (K=50)
Select 2D (Reference 2D)
Filament Tracer (with 2D reference)
Inspect Pick
Extract Particles (2.38 Å/pix)
462,672 particles
2D Class (K=50)
Select 2D
Extract Particles (1.55 Å/pix)
Helical Refinement 4.96 Å
(Sym: C5)
Symmetry Search
(Rise 5-50 / Twist 5-50)
Extract Particles (1.55 Å/pix)
201,039 particles
Helical refinement 3.32 Å
(Sym: C5)
Reference-Based Motion Corr.
Helical Refinement 3.44 Å
(Sym: C5, Rise: 12.26, Twist 29.00° )
Local CTF Refinement
Helical Refinement
2 Rounds
GSFSC Resolution: 3.11Å
1.0
0.8
0.6
0.4
0.2
0.0
DC
15Å
7.6Å
5.1Å
3.8Å
3.1Å
2.5Å
No Mask (3.5Å)
Spherical (3.3Å)
Loose (3.2Å)
Tight (3.1Å)
Corrected (3.1Å)
Sym: C5, Rise: 12.26 Å, Twist: 28.96°

D

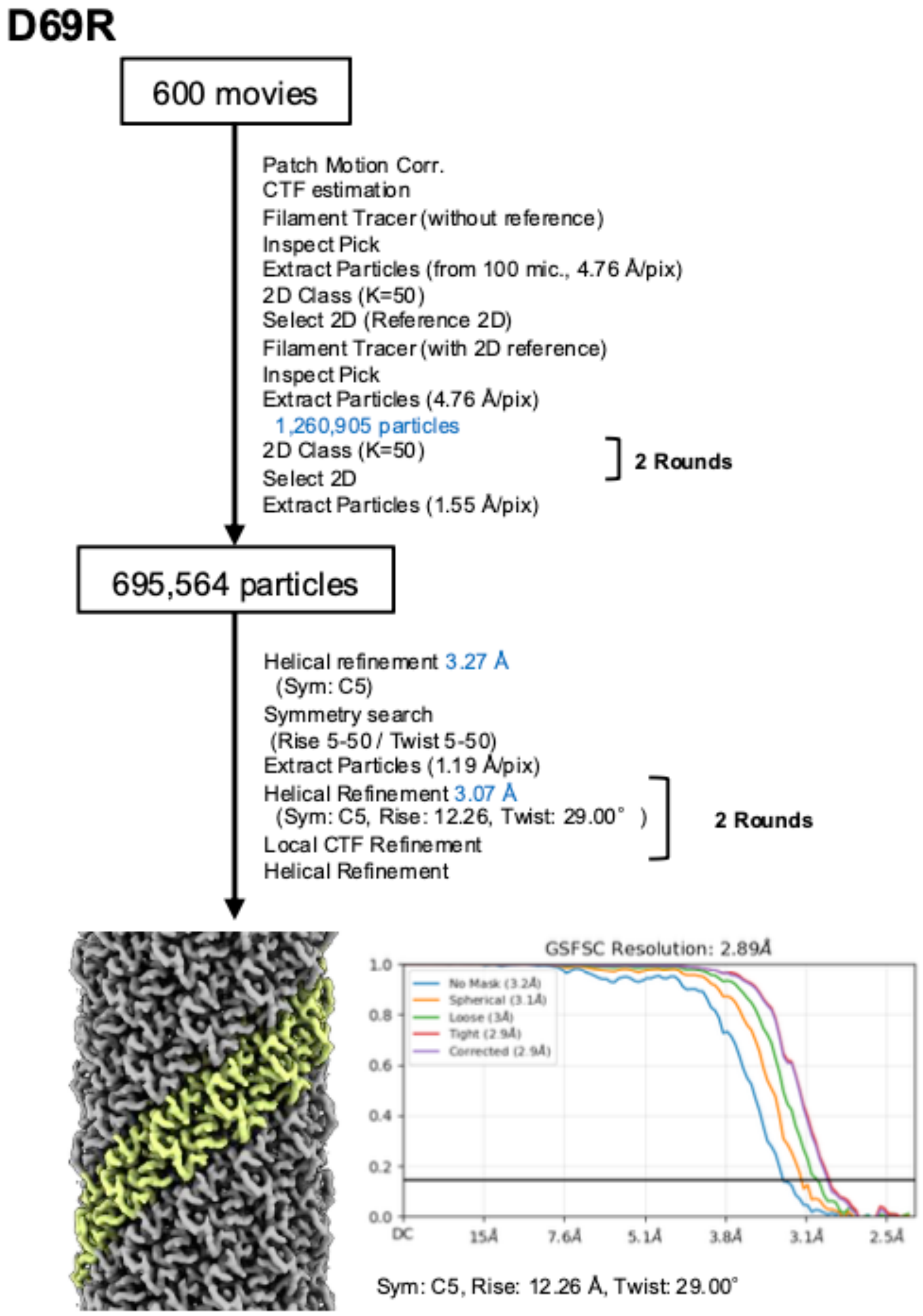


**Supplementary Figure 1**. Cryo-EM data processing workflow for the H pilus mutants. All the processing was performed in cryoSPARC (v.4.6). Gold-standard FSC curve of the final map was shown. The resolution cut-off was at FSC=0.143.

A

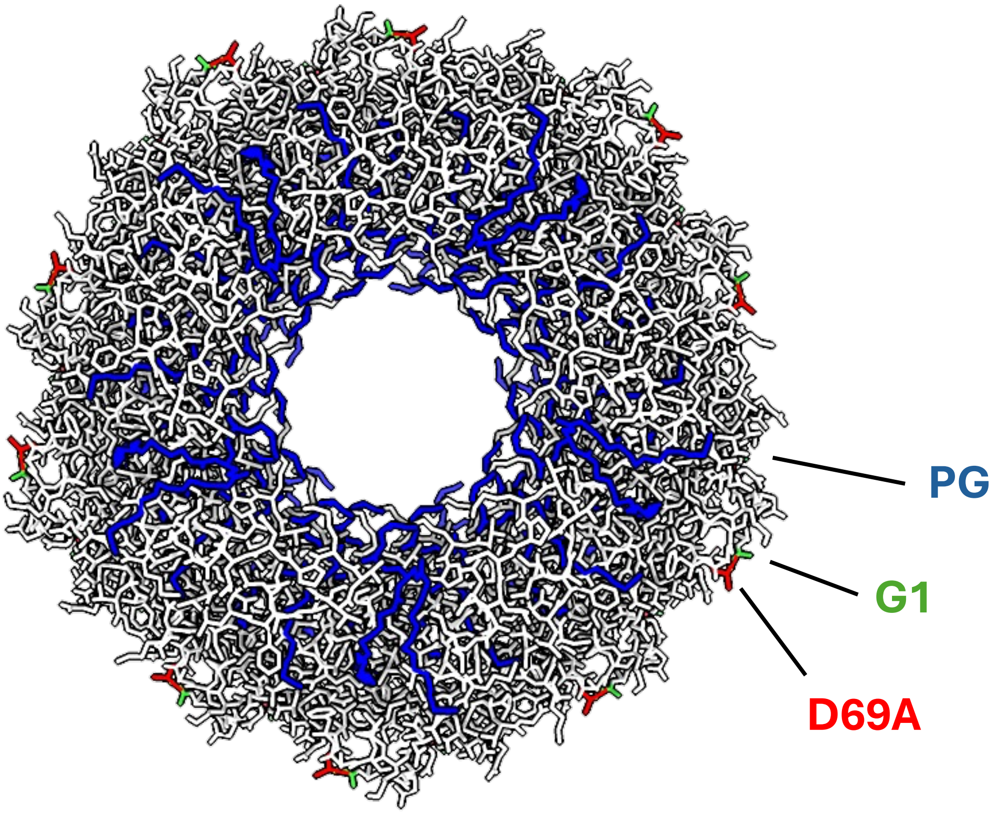


B

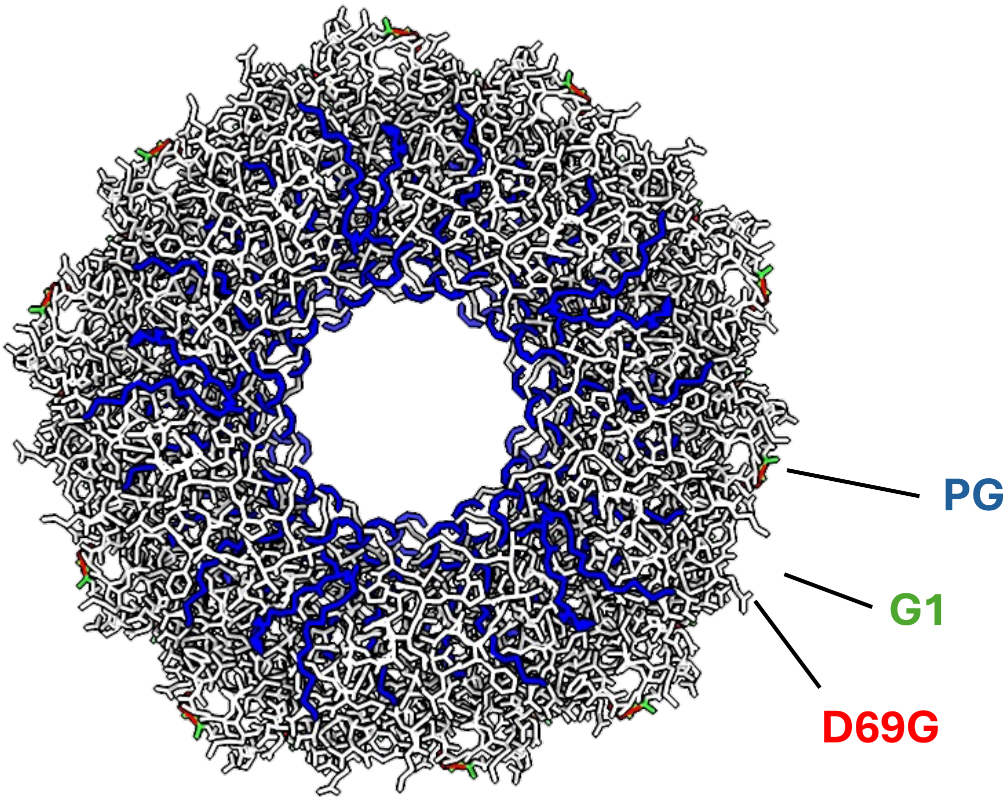

**C**

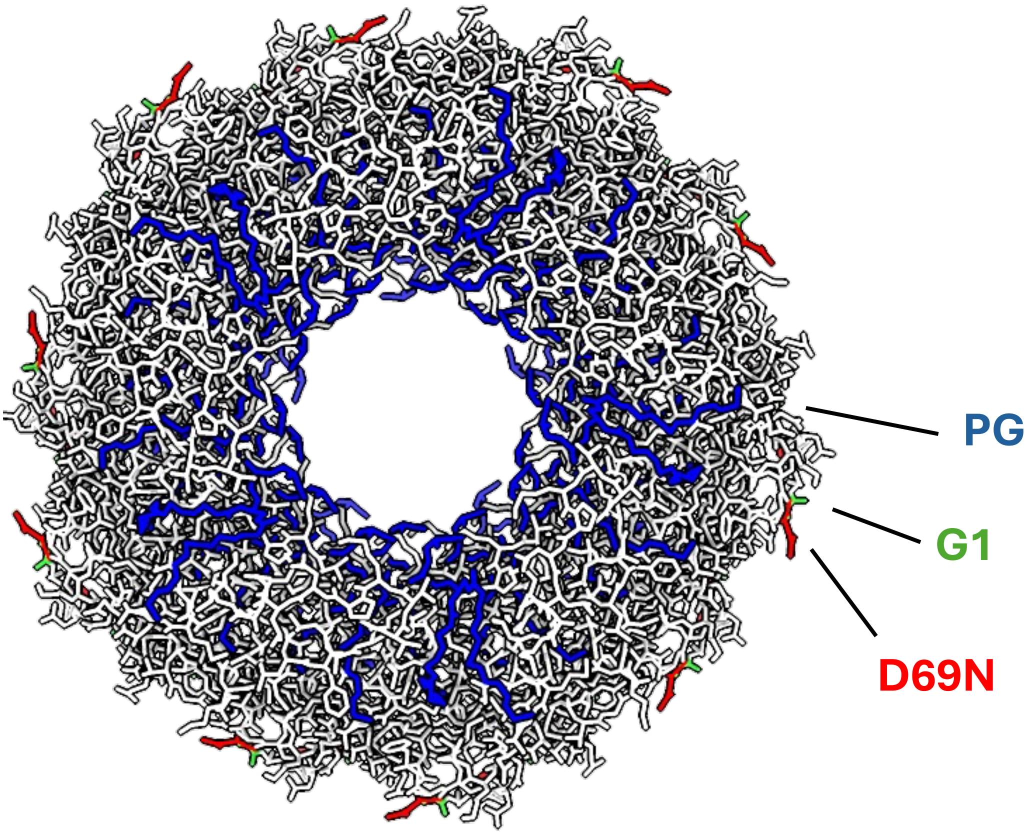


**Supplementary Figure 2.** Mapping of the ThrA mutant side chains on the surface of the assembled pilus. Colour scheme as in Figure 4.

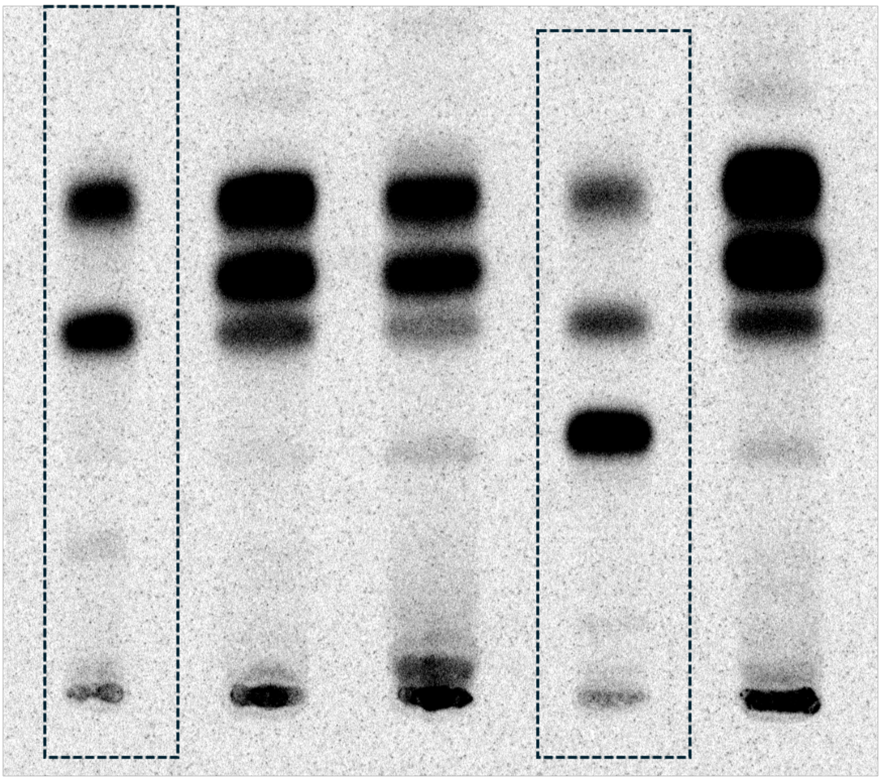

**Supplementary Figure 3.** Uncropped TLC related to Fig 5.

**Table S1.** Primers used in this study.

| Primers | Sequences | Description |
|---|---|---|
| F_WT_trhA | GTTATCCCTAAGTGACTTTTACGATAAGC | Amplifies the *trhA* gene from R27 plasmid |
| R_WT_trhA | CTCTATTCTTGATGGAGCTAGGGATAAC | |
| F_D69_SDM | GCTGGTATTCCTCTGTGA | Amplifies liner vector pSEVA612s for all the D69X mutagenesis |
| R_69N_SDM | GTTCAGGAACGAGCTGA | Amplifies liner vector pSEVA612s for the D69N mutagenesis |
| R_69R_SDM | GCGCAGGAACGAGCTGA | Amplifies liner vector pSEVA612s for the D69R mutagenesis |
| R_D69A_SDM | CGCCAGGAACGAGCTG | Amplifies liner vector pSEVA612s for the D69A mutagenesis |
| R_D69G_SDM | GCCCAGGAACGAGCTGATA | Amplifies liner vector pSEVA612s for the D69G mutagenesis |
| F_D69K_SDM | CTCGTTCCTGaaaGCTGGTATTC | Amplifies liner vector pSEVA612s for the D69K mutagenesis |
| F_G1S_SDM | TGCGTACGCGagcTCCGATGATG | Amplifies liner vector pSEVA612s for the G1S mutagenesis |
| R_G1S_SDM | AAACTGCAATTTGCTAAAACCAAGAG | |
| F_G1K_SDM | TGCGTACGCGaaaTCCGATGATG | Amplifies liner vector pSEVA612s for the G1K mutagenesis |
| R_G1K_SDM | AAACTGCAATTTGCTAAAAC | |
| F_G1D_SDM | TGCGTACGCGgatTCCGATGATG | Amplifies liner vector pSEVA612s for the G1D mutagenesis |
| R_G1D_SDM | AAACTGCAATTTGCTAAAACCAAGAGAATAAAC | |
| F_IncH pilin seq | GCTCCATCAAGAATAGAGG | Sanger sequencing primer to check the amino acid substitution |
| F_IncH pilin seq | CCAATGCGTTTTCGTGA | |

**Table S2.** Cloning plasmids used for mutagenesis.

| **Vector** | **Description** |
|---|---|
| pACBSR | SmR; expresses I-SceI and lambda-red induced by L-Ara. |
| pSEVA612S | GmR; integrative plasmid (ori R6K) that harbours the oriT for tri-parental mating. |
| pSEVA612S_*trhA* WT | pSEVA612S derivative; used for the building the site-direct mutagenesis constracts for *trhA* from R27 |
| pSEVA612S_Δ *trhA* | pSEVA612S derivative; used for the deletion of *trhA* gene from R27 |
| pSEVA612S_*trhA* D69A | pSEVA612S derivative; used for amino acid substitution of D69A from TrhA |
| pSEVA612S_*trhA* D69R | pSEVA612S derivative; used for amino acid substitution of D69R from TrhA |
| pSEVA612S_*trhA* D69G | pSEVA612S derivative; used for amino acid substitution of D69G from TrhA |
| pSEVA612S_*trhA* D69N | pSEVA612S derivative; used for amino acid substitution of D69N from TrhA |
| pSEVA612S_*trhA* D69K | pSEVA612S derivative; used for amino acid substitution of D69K from TrhA |
| pSEVA612S_*trhA* G1K | pSEVA612S derivative; used for amino acid substitution of G1K from TrhA |

**Table S3.** Conjugative plasmids and strains used and generated in this study.

| **Plasmids** | **Description** |
|---|---|
| R27 | Prototype plasmid for IncH, and it is a derepressed plasmid. Its cyclic region is 67F-74L. |
| R27 Δ67-74 | R27 plasmid with a deletion of TrhA amino acid from 67-74, considers as no conjugative pilus production. |
| R27 D69A | R27 plasmid with an amino acid substitution of TrhA D69A |
| R27 D69N | R27 plasmid with an amino acid substitution of TrhA D69N |
| R27 D69G | R27 plasmid with an amino acid substitution of TrhA D69G |
| R27 D69R | R27 plasmid with an amino acid substitution of TrhA D69R |
| R27 D69K | R27 plasmid with an amino acid substitution of TrhA D69K |
| R27 G1K | R27 plasmid with an amino acid substitution of TrhA G1K |

| **Strains** | **Description** |
|---|---|
| CC118 λ pir | Expresses the Pi protein for the replication of plasmids with the R6K origin. |
| *E. coli* 1047 pRK2013 | Triparental conjugation helper strain. Kanamycin resistant |
| *E. coli* | *E. coli* K-12 strain |
| *E. coli -trp* | Donor strain for conjugation assays. It is *trp*- mutant, won't be able to grow on the minimal media. |
| *K. pneumoniae* | ICC8001. Parental wild type (WT) strain of K. pneumoniae ATCC43816 serially passaged in vitro on Rifampicin(100µg/ml) followed by two passages in BALB/c mice. |
| *E. cloacae* | *Enterobacter cloacae* ATCC13047 |
| *C. amalonaticus* | CMS57 isolate 57.41.1 / Citrobacter.amalonaticusC3H ICC3000 |
| EPEC | *Enteropathogenic E. coli* e2348/69 |
| AL95 | A *E. coli* strain lacking phosphatidylethanolamine; Δ*pssA* |
| W3110 | The wild type *E. coli* strain for AL95 |

**Table S4.** Data collection, processing and refinement statistics for the H pilus mutants.

| | D69N | D69A | D69G | D69R |
|---|---|---|---|---|
| **PDB entry** | 9VP3 | 9VP2 | 9VPE | 9VP4 |
| **EMDB entry** | EMD-65235 | EMD-65234 | EMD-65250 | EMD-65236 |
| **Data collection and processing** | | | | |
| Magnification | 120 000 | | | |
| Microscope | Glacios 2 | | | |
| Voltage (kV) | 200 | | | |
| Detector | Falcon 4 | | | |
| Electron Dose ($e^-$/Å) | 46 | 48 | 46 | 46 |
| Deforcus range (µm) | -0.8 to -1.8 | | | |
| Pixel size (Å) | 1 192 | | | |
| Data Processing Program | cryoSPARC (v.4.6.1) | | | |
| Movies | 600 | 711 | 200 | 600 |
| Initial / Final particle images (n | 1,834,272 / 1,477,420 | 1,256,631 / 420,903 | 462,670 / 200,984 | 1,260,905 / 695,564 |
| Symmetry imposed | C5 | | | |
| Helical rise (Å) | 12 24 | 12 24 | 12 26 | 12 26 |
| Helical twist (°) | 29 00 | 29 00 | 28 96 | 29 00 |
| Map resolution (Å) | 2 88 | 3 15 | 3 11 | 2 89 |
| FSC threshold | 0 143 | 0 143 | 0 143 | 0 143 |
| **Refinement** | | | | |
| Refinement Program | PHENIX (v.1.20.1) | PHENIX (v.1.20.1) | PHENIX (v.1.20.1) | PHENIX (v.1.20.1) |
| Model resolution (Å) | | | | |
| FSC threshold = 0.143 | 2 82 | 3 13 | 3 04 | 2 85 |
| R.m.s. deviations | | | | |
| Bond length (Å) | 0 002 | 0 003 | 0 002 | 0 002 |
| Bond angles (°) | 0 388 | 0 412 | 0 333 | 0 408 |
| Validation | | | | |
| MolProbity score | 1 16 | 1 05 | 0 94 | 1 44 |
| Clashscore | 1 77 | 2 67 | 1 79 | 4 40 |
| Ramachandran plot | | | | |
| Favored / Allowed (%) | 100 / 0 | 100 / 0 | 100 / 0 | 100 / 0 |
| Disallowed (%) | 0 00 | 0 00 | 0 00 | 0 00 |
| Mask CC | 0 85 | 0 87 | 0 88 | 0 87 |
